# Astrocytic CREB in nucleus accumbens promotes susceptibility to chronic stress

**DOI:** 10.1101/2024.01.15.575728

**Authors:** Leanne M. Holt, Trevonn M Gyles, Eric M. Parise, Angelica Minier-Toribio, Tamara Markovic, Matthew Rivera, Szu-Ying Yeh, Eric J. Nestler

## Abstract

**Background:** Increasing evidence implicates astrocytes in stress and depression in both rodent models and human Major Depressive Disorder (MDD). Despite this, little is known about the transcriptional responses to stress of astrocytes within the nucleus accumbens (NAc), a key brain reward region, and their influence on behavioral outcomes.

**Methods:** We used whole cell sorting, RNA-sequencing, and bioinformatic analyses to investigate the NAc astrocyte transcriptome in male mice in response to chronic social defeat stress (CSDS). Immunohistochemistry was used to determine stress-induced changes in astrocytic CREB within the NAc. Finally, astrocytic regulation of depression-like behavior was investigated using viral-mediated manipulation of CREB in combination with CSDS.

**Results:** We found a robust transcriptional response in NAc astrocytes to CSDS in stressed mice, with changes seen in both stress-susceptible and stress-resilient animals. Bioinformatic analysis revealed CREB, a transcription factor widely studied in neurons, as one of the top-predicted upstream regulators of the NAc astrocyte transcriptome, with opposite activation states seen in resilient versus susceptible mice. This bioinformatic result was confirmed at the protein level with immunohistochemistry. Viral overexpression of CREB selectively in NAc astrocytes promoted susceptibility to chronic stress.

**Conclusions:** Together, our data demonstrate that the astrocyte transcriptome responds robustly to CSDS and, for the first time, that transcriptional regulation in astrocytes contributes to depressive-like behaviors. A better understanding of transcriptional regulation in astrocytes may reveal unknown molecular mechanisms underlying neuropsychiatric disorders.

## Introduction

Stress-related disorders remain one of the world’s greatest public health burdens. To date, research has focused primarily on the underlying persistent molecular mechanisms contributing to neuronal dysfunction within limbic regions of the brain, including transcriptional and epigenetic modifications mediating such lasting changes. However, emerging evidence indicates that non-neuronal cells, including astrocytes, may play a larger role than previously believed [1–3]. Astrocytes are well known for their ability to regulate synapse formation and function, neuronal transmission, blood brain barrier, and metabolic coupling—all biological processes implicated in stress and depression [1–3]. Indeed, changes in astrocyte morphology, number, and function have been observed in both rodent stress models and postmortem studies of human depression across many brain regions [4–12]. Antidepressant treatments, including selective serotonin reuptake inhibitor (SSRIs) and ketamine, result in alterations to astrocytes, including reversal of stress-induced changes in astrocyte morphology [13–16]. Furthermore, manipulating astrocyte function bidirectionally influences depressive-like behaviors in rodent models, demonstrating the active role of astrocytes in mediating complex behavior, including those associated with stress and depression [6, 8, 11, 17–21].

Previous work using targeted molecular approaches demonstrates dysregulation of astrocytic gene expression in rodent stress models and in human Major Depressive Disorder (MDD), and emerging evidence from unbiased RNA-sequencing (RNA-seq) studies of bulk tissue and single cells across several brain regions also implicate astrocytes [9, 12, 22–26]. However, astrocyte-specific RNA-seq in rodents – across disease models – has only been performed in the prefrontal cortex (PFC), hippocampus, and amygdala [9, 24, 25, 27, 28]. Furthermore, while there has been an increase in sequencing studies focused on astrocytes, we know surprisingly little regarding transcriptional regulators in astrocytes. Within the nucleus accumbens (NAc), a region important for motivation, reward, and learning, and heavily implicated in stress and depression, alterations in astrocyte number, morphology, and synapse association have been observed [1, 29–31]. Despite this early literature, transcriptional responses of NAc astrocytes and their potential role in regulating behavioral consequences of chronic stress remain unknown.

We performed RNA-seq on whole-cell sorted NAc astrocytes following chronic social defeat stress (CSDS), a highly validated procedure used to study depression and other human stress disorders [1]. Bioinformatic analysis revealed a robust transcriptional response in astrocytes from both resilient and susceptible animals. Additional analysis identified key predicted astrocytic transcriptional regulators, including the transcription factor, cAMP Response Element-Binding Protein (CREB). Transcriptional regulation of neuronal CREB has been extensively studied in neuropsychiatric disorders, with increased CREB activity within NAc neurons associated with depressive behaviors [31, 32]. While CREB has been implicated as a transcriptional regulator in cultured astrocytes, its role in astrocytes *in vivo*, including any effects on behavior, remain elusive [33–36]. We show that viral manipulation of CREB selectively in NAc astrocytes resulted in a bias towards susceptibility to CSDS. In summary, we demonstrate, for the first time, that a transcription factor in NAc astrocytes controls behavioral response to stress.

## Methods

### Animals

All experiments were performed with wildtype C57BL/6 male mice (aged 9–10 weeks, The Jackson Laboratory) according to NIH guidelines and with approval from the Animal Care and Use Committee of Mount Sinai. All animals were group-housed and maintained on a normal 12 hr light/dark cycle (07:00 lights on; 19:00 lights off) with food and water available *ad libitum*. Immediately prior to and during CSDS, mice were single-housed. Every effort was made to minimize pain and discomfort.

### Chronic Social Defeat Stress (CSDS) + Social Interaction (SI) testing

Experimental C57BL/6 mice are introduced into the cage of resident CD1 retired breeder mice pre-screened for aggression [37, 38]. This is repeated for 5 minutes daily over 10 days, with housing across a perforated Plexiglass divider for the remaining 24 hr to allow sensory exposure. Control mice are housed with a Plexiglass divider between another wildtype control mouse and rotated to a different cage daily. The SI test for social avoidance behavior was performed within 24 hr of the last CSDS session. Animal exploratory behavior was recorded in an open-field arena containing a wire enclosure. This enclosure was empty for the first 2.5 mins (no target present) and subsequently held a novel CD1 aggressor in the second 2.5 mins (target present). The experiment was conducted under red light conditions. Social avoidance (Social Interaction Ratio) was calculated by dividing the time spent in the interaction zone when the target mouse was present over the time spent in the zone when the target mouse was absent. Using the SI ratio, mice were categorized as either susceptible (SI ratio < 0.9) or resilient (SI ratio > 1.1), a measure that has been shown to be highly predictive of numerous other behavioral sequelae after CSDS [37, 38].

### Astrocyte Isolation

Astrocytes were isolated using Miltenyi BioTech’s ACSA-2 MicroBead Kit as previously described [39, 40] from freshly micropunched NAc tissue (bilateral 14G punches). Following astrocyte elution, a final 300 rcf centrifuge spin was performed, the supernatant removed, and astrocytes resuspended in 300 µL Tri-Reagent (Zymo). To ensure lysis prior to snap-freezing on dry ice, the resuspension was vortexed for 30 seconds. All samples were stored at -80°C until RNA extraction.

### RNA Extraction, Library Preparation, and Sequencing

RNA was extracted using Zymo’s Directzol RNA MicroPrep kit following manufacturer’s instructions. RNA quality and quantity was assessed using Bioanalyzer (Agilent). Any sample with a RIN value of less than 7.8 was excluded. 500 pg of RNA was used as input in Takara’s SMARTer® Stranded Total RNA-Seq Kit v2 – Pico, with ribodepletion, according to the manufacturer’s instructions. Sequencing libraries were generated for each sample individually using Takara’s Unique Dual Index Kit. Following library preparation, sequencing was performed with Genewiz/Azenta on an Illumina Novaseq with a 2×150 bp paired-end read configuration to produce 40M reads per sample. Quality control was performed using FastQC. All raw sequencing reads underwent adapter trimming and were mapped to mm10. Genes with a row sum less than 10 were excluded prior to differential gene expression analysis. Differential expression was performed in R version 4.0.2 using the DESeq2 package version 1.28.1 25516281, with built-in independent filtering disabled. For differential expression analysis, Resilient (n = 7) and Susceptible (n = 7) astrocytes were compared to Control astrocytes (n = 8). Significant DEGs were determined by a 20% Log2FC and p < 0.05. Volcano plots were generated in R (v4.0.2) using tidyverse package (v1.3.1) and ggplot (v3.4.2). Venn diagrams were generated using nVenn [41]. Union heatmaps were generated using Morpheus (https://software.broadinstitute.org/morpheus).

### Rank Rank Hypergeometric Overlap (RRHO)

RRHO plots were generated using the RRHO2 package (github.com/RRHO2/RRHO2). Human RNA-seq data from the NAc was accessed from [42].

### Gene Ontology analysis

Gene ontology for “Biological Processes 2021” database was performed in R using the enrichR package with our filtered DEG lists as input [43, 44]. Plots were made with ggplot2 (v3.4.2). Specific terms presented are summarized if redundancies were present.

### Ingenuity upstream regulator analysis

Predicted upstream regulators were identified using Ingenuity Pathway Analysis software (Qiagen) with the identified DEGs lists as input. Upstream regulators were filtered to only include those considered as “molecules” with Benjamini-Hochberg corrected p-values for p < 0.05 and an activation z-score of greater or lesser than 0.2 [45, 46].

### Immunohistochemistry

Brains were collected within 48 hr of the SI test. At time of collection, animals were deeply anesthetized with peritoneal injections of 500mg/kg Fatal Plus and intracardially perfused with PBS, followed by 20 mL 4% paraformaldehyde (PFA). Brains were post-fixed for 72 hr, and subsequently sliced at 30uM sections. Sections were blocked for 1 hr in blocking buffer, followed by overnight incubation with primary antibodies in diluted blocking buffer (1:3 of blocking buffer in PBS). Sections were then washed for 15 mins (3x) in diluted blocking buffer before being incubated with secondary antibodies (all 1:500) in diluted blocking buffer for 1.5 hr at room temperature in the dark. Three additional 15 min washes in diluted blocking buffer followed the incubation in secondary antibodies (all 1:500). Finally, sections were counterstained with DAPI (1:10,000) for five minutes before a final wash in PBS. Sections were mounted with Fluoromount medium (Sigma #F4680) on glass slides (FisherScientific #12-550-15) and covered with cover glass (FisherScientific #12-548-5E). The following primary antibodies were used: NeuN (MilliporeSigma #ABN91; 1:1000), Sox9 (Abcam # ab76997; 1:500), total CREB (Cell Signaling #9197L; 1:500), pCREB (Cell Signaling #9198S; 1:500). The following secondary antibodies were used: Ch-488 (Jackson #703-546-155), Ms-Cy3 (#715-166-150), Rb-647 (#711-606-152), Rb-Cy3 (#711-166-152).

### Confocal microscopy

Images were taken on a Zeiss LSM780 microscope with a 40x oil immersion lens. For quantification of CREB and pCREB, the settings for laser power and gain were determined for control samples and subsequently not adjusted. The experimenter was blinded to imaging for the Res and Sus samples. Integrated intensity of CREB and pCREB were determined using Cell Profiler.

### AAV viruses

Control EGFP (pAAV.GfaABC1D.PI.Lck-GFP.SV40; #105598-AAV5) and tdTomato (pZac2.1 gfaABC1D-tdTomato; #44332-AAV5) viruses were purchased from Addgene [47]. Astrocyte-specific CREB viruses were generated by Virovek by cloning the GFP-CREB and GFP-mCREB sequence [48–50] with the GfaABC1D promoter into an AAV2/5.

### Stereotaxic surgery

Mice were anesthetized with a mixture of ketamine (100mg/kg) and xylazine (10mg/kg) and positioned in a small-animal stereotactic instrument (Kopf Instruments). The skull surface was exposed, and 33-g syringe needles (Hamilton) used to bilaterally infuse 1µL of AAVs (diluted in sterile PBS to 2 x 10^10^) at a rate of 0.2µL/min. NAc coordinates relative to the Bregma were: AP +1.3, ML +1.5, DV -4.4 at a 10° angle.

### Statistical analyses

All data were plotted as mean +/- SEM, and statistical analysis was performed in Prism 10. Behavioral data was analyzed using one-way or two-way ANOVAs, as appropriate, followed by Tukey’s post-hoc test. Immunohistochemistry data was analyzed using one-way ANOVA followed by Tukey’s post-hoc test and Pearson r correlation. Outliers were determined for immunohistochemistry data using Prism’s ROUT method, with the strictest cutoff of Q = 0.1%.

## Results

### Astrocyte transcriptome robustly responds to chronic stress

To better understand the role of astrocytes in stress and depression, we performed astrocyte whole-cell sorting and RNA-seq after ten days of CSDS (Fig 1A). Male C57BL/6 mice were exposed to different CD1 aggressors for 5 minutes every day for 10 consecutive days with overnight sensory exposure (plexiglass separators in cage). Within 24 hr of the last defeat session, an SI test was performed and the animals were subsequently separated into resilient (Res) and susceptible (Sus) categories as previously published [37, 38]. Animals that displayed a high interaction ratio and interaction time, and low time in the corners, were considered Res; animals with a low interaction ratio and interaction time, and high time in the corners, were considered Sus (Fig 1B-D). Within 48–72 hr of the last defeat session, astrocytes were collected from the NAc with stress categories counterbalanced across collection days and then RNA-seq was performed.

**Figure 1.**
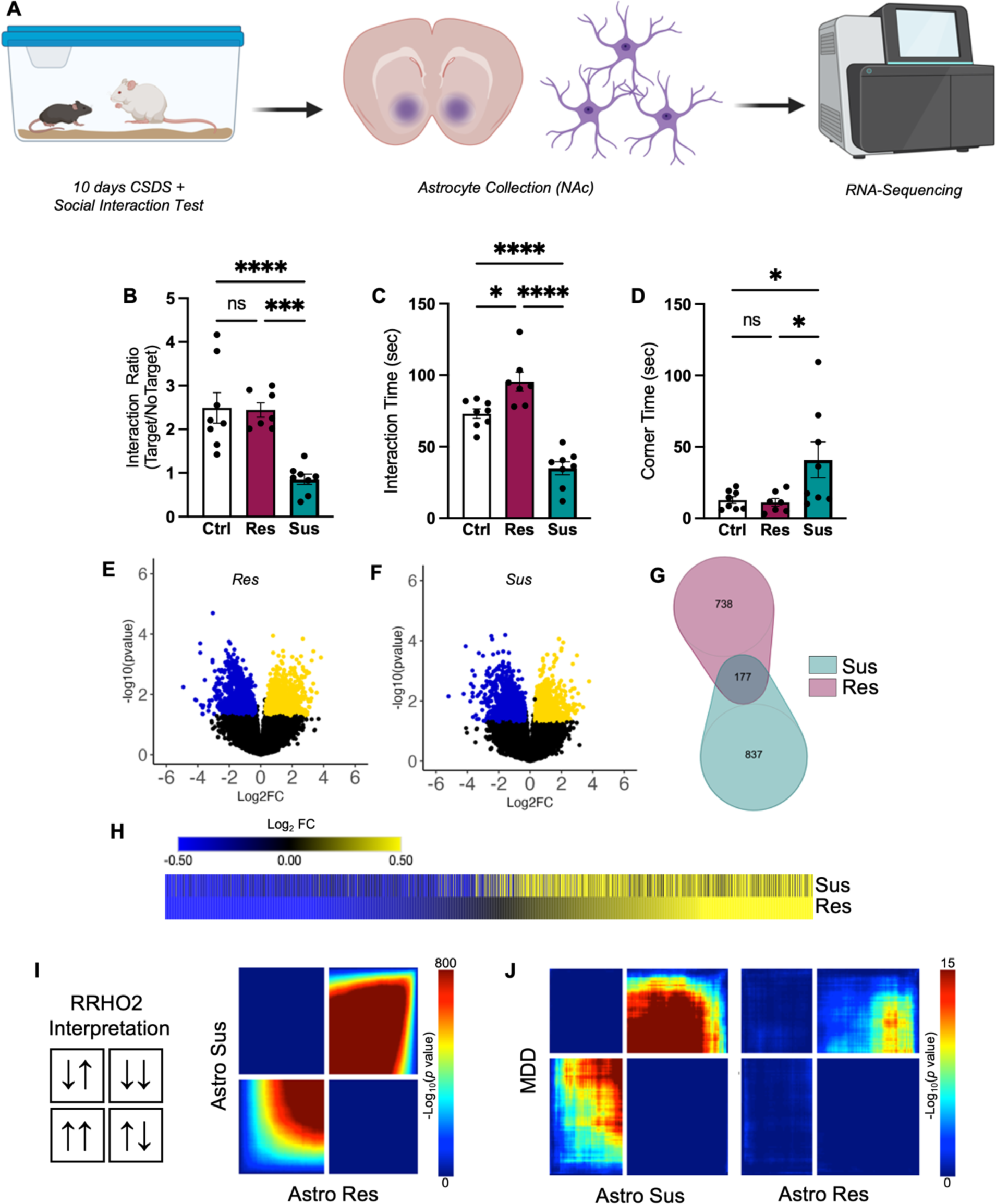
Astrocyte transcriptome in NAc responds robustly to chronic stress. A) Cartoon illustration of experimental design. B–C) Behavior of phenotyped animals selected for astrocyte collection and RNA-seq. CSDS resulted in Res (purple) and Sus (teal) phenotypes, as expected. B) Sus animals demonstrate decreased Interaction Ratios compared to Ctrl and Res animals (F_(2,20)_ = 15.63, *P* < 0.0001; Tukey post-hoc \*\*\**p* < 0.001). C) Res animals demonstrate an increase in time interacting with a novel CD1 aggressor compared to Ctrl and Sus animals, while Sus animals demonstrate a decrease compared to Ctrl animals (F_(2,20)_ = 38.44, *P* < 0.001; Tukey post-hoc *p < 0.05; ****p < 0.0001). D) In contrast, Sus animals demonstrate an increase in time spent in corners compared to both Res and Sus (F_(2,20)_ = 4.636, *P* = 0.0222; Tukey post-hoc *p < 0.05). Volcano plots of detected genes from E) Res and F) Sus astrocytes compared to Ctrl. Upregulated DEGs are indicated in yellow, while downregulated DEGs are indicated in blue. G) Venn diagram reveals little overlap between significant DEGs in Res (purple) and Sus (teal) astrocytes. However, H) union heatmap demonstrates considerably similar Log_2_FC expression of significant DEGs where again upregulated DEGs are indicated in yellow and downregulated DEGs are indicated in blue. The union heatmap nevertheless does highlight DEGs that are regulated differently in Res versus Sus. I) RRHO2 threshold-free genome-wide comparison confirms the union heatmap by revealing partly concordant gene expression between Res and Sus. J) RRHO2 comparison of human MDD bulk RNA-seq of NAc to Sus (left) and Res (right) RNA-seq of NAc astrocytes demonstrates some concordant expression between the mouse and human datasets. Data represented at mean +/- SEM, n = 7–8 animals per condition. RNA-seq: RNA-sequencing; CSDS: chronic social defeat stress; Res: resilient; Sus: susceptible; Ctrl: control; DEG: differentially expressed genes; Log_2_FC: log2 fold change; RRHO2: rank rank hypergeometric overlap.

Volcano plots from the differential expression analysis (DESeq2) revealed a robust transcriptional response in astrocytes obtained from both Res and Sus animals in comparison to control astrocytes, with an equal distribution across up- and downregulated genes (Fig 1E, F). The venn diagram comparison of the identified statistically-significant, differentially expressed genes (DEGs; Fig 1G) revealed little overlap (11%, 177 DEGs) between Res and Sus astrocytes. However, union heatmaps of DEGs revealed generally similar patterns of gene expression (Fig 1H). Genes downregulated in Res astrocytes were largely also downregulated in Sus astrocytes, with some exceptions. There appeared to be less convergence in upregulated DEGs between Sus and Res astrocytes. We also observed that, overall, the magnitude of the Log_2_FC was greater in Res compared to Sus astrocytes (Fig 1E-F).

To determine if this pattern of gene expression was confined to those genes that were determined to be statistically significantly affected by CSDS or were found genome-wide, we utilized Rank Rank Hypergeometric Overlap (RRHO2). RRHO2 compares changes in gene expression between two datasets in a “threshold-free” manner [51]. RRHO2 revealed, similar to our union heatmaps, that CSDS induced some concordant gene expression changes in Res and Sus astrocytes from the NAc (Fig 1I). This finding suggests that, while distinct genes are found in Res and Sus astrocyte populations with statistical thresholds, there is also a large population of dysregulated genes in astrocytes associated with a general stress response.

Finally, given that bulk tissue and single-cell/nuclei RNA-seq has implicated astrocyte transcriptomic dysfunction in tissue from human MDD patients, we determined if astrocytes from our rodent CSDS model exhibited similar transcriptional patterns compared to the human MDD transcriptome. We compared our astrocyte RNA-seq data to available human bulk RNA-seq using RRHO2. We found concordance between Sus and Res astrocytes, albeit to a lesser extent in Res astrocytes, compared to human MDD patients (Fig 1J).

To determine the molecular, downstream impact of CSDS on the astrocyte transcriptome we performed Gene Ontology (GO) analysis on the Res and Sus astrocyte DEGs. We utilized EnrichR GO: Biological Pathways 2021 [43, 44] and identified more significant GO Terms in Res compared to Sus astrocytes (111 vs 52), with only three overlapping Terms (Fig 2A). This is perhaps not surprising, as while global gene expression appears to be similar between Res and Sus astrocytes, there are only a few overlapping significant DEGs between Res and Sus. The top 10 GO terms implicated in Res astrocytes were: cytoplasmic translation (*cytoplasmic sequestering of NF-kappaB*; *cytoplasmic translation*), neuronal dysfunction (*axon ensheathment in central nervous system*), and cytoskeletal integrity (*microtubule depolymerization; microtubule polymerization or depolymerization*) (Fig 2B). In contrast, post-translational protein stability (*protein targeting, establishment of protein localization to membrane; protein palmitoylation*) and cellular processing (*cellular response to peptide hormone stimulus; copper ion transport*) were implicated in Sus astrocytes (Fig 2C).

**Figure 2.**
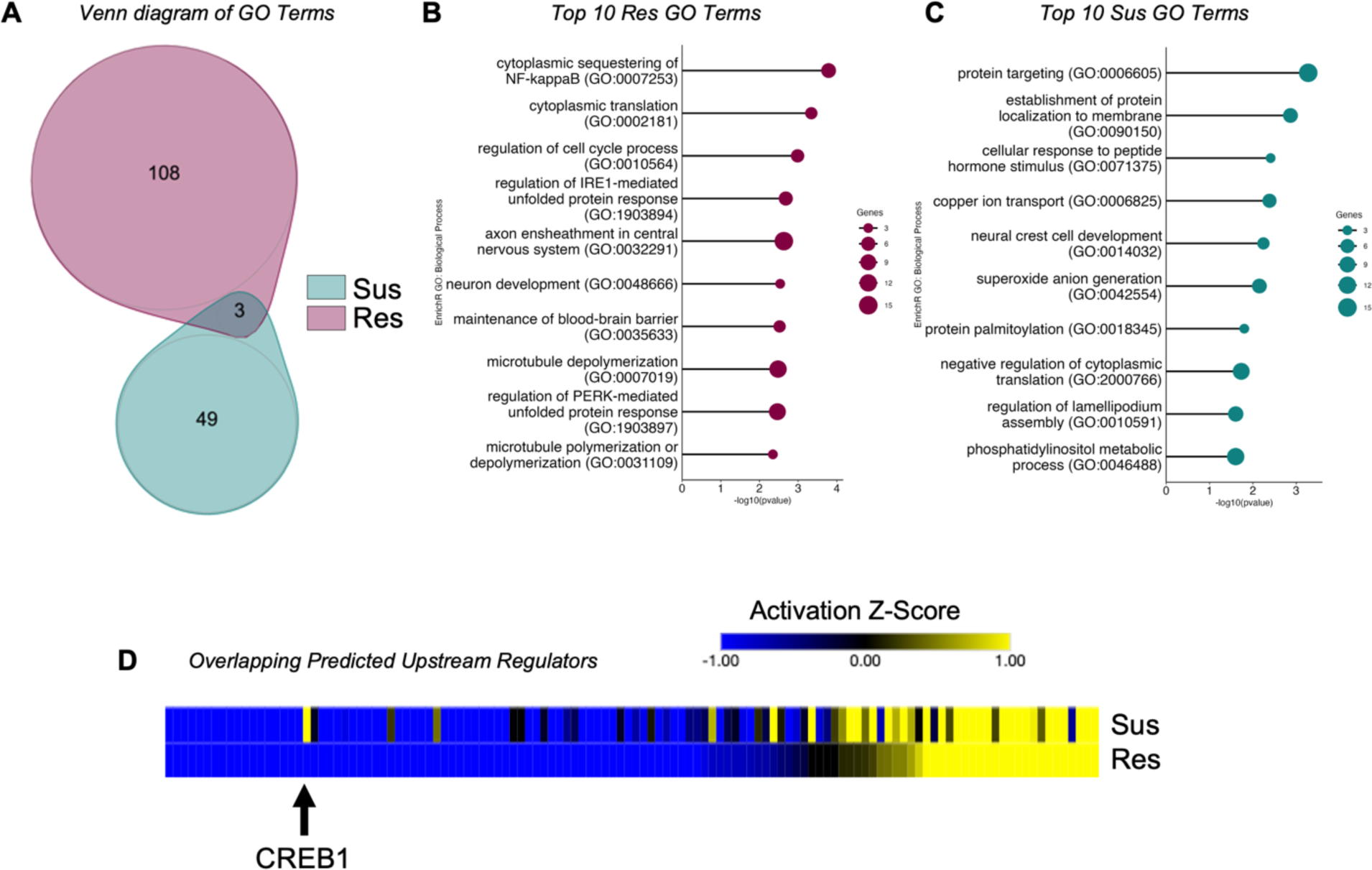
The astrocyte transcriptomic response to stress in NAc is phenotypically specific. A) Venn diagram of identified GO terms between Res (purple) and Sus (teal) astrocytes reveals little overlap of downstream molecular consequences of stress on astrocytes. The top 10 identified GO Terms of B) Res and C) Sus further implicate distinct molecular responses between the two phenotypes. D) Union heatmap of overlapping upstream regulators reveals similar activation states in the majority, while several demonstrate opposing states of activation (blue indicates inhibited regulators, yellow indicates activated regulators). The black arrow demarcates CREB1, which demonstrates a predicted activation z-score in Sus and inhibition in Res astrocytes. Res: resilient; Sus: susceptible.

We next determined potential transcriptional mediators governing the astrocyte transcriptome in response to chronic stress by performing Ingenuity Pathway Analysis’s Upstream Regulator analysis. More upstream regulators were identified in Res compared to Sus astrocytes (270 vs. 181), with only 124 in common between Res and Sus astrocytes (Fig 2D). We identified several upstream regulators previously implicated in social behavior, stress, or depression, including CD38, TCF7L2, VEGF, and BDNF [37, 52–55]. Of the overlapping regulators, we found that the majority were activated or inhibited (direction of activation Z-score in Fig 2D) similarly in both Res and Sus (80%, 99/124), with only 25 (20%) found to be incongruent (Fig 2D). This again highlights both an overall general transcriptional response to chronic stress in astrocytes, as well as potential transcriptional regulators that may govern differential molecular and behavioral consequences.

### Astrocytic CREB is regulated by chronic stress

CREB1 caught our attention (Fig 2D, arrow) in the upstream regulator analysis, as it was strongly predicted in both Res and Sus astrocytes, but with opposing directions in the activation z-score. CREB is a well-known transcription factor that influences a variety of neuropsychiatric disorders, including stress and depression in both rodent models and human MDD patients [31, 32] Previous work has implicated CREB as a transcriptional regulator in astrocytes [33, 34], however, the role of astrocytic CREB in stress and depression has yet to be investigated. We first validated our bioinformatic findings to determine if CREB was indeed activated in NAc astrocytes in response to CSDS. Male C57B/L6 mice were exposed to our 10-day CSDS paradigm, with SI testing to determine Res and Sus animals followed by perfusion and tissue collection. SOX9 immunohistochemistry was used to demarcate astrocytes in combination with antibodies for total CREB or phosphorylated CREB (pCREB) (Fig 3A,B). Confocal microscopy revealed a significant reduction in total CREB levels in Sus astrocytes of NAc compared to both Ctrl and Res astrocytes with no significant difference between Res and Ctrl. The effect seen in Sus astrocytes was observed when both subregions of the NAc were considered separately (Core: Fig 3D; Shell: Fig 3E). Importantly we observed a statistically significant positive correlation between SI Ratio and mean CREB integrated intensity (analyzing the entire NAc) for individual animals, demonstrating that the change in astrocytic CREB is not dependent on our categorical Res and Sus assignments (Fig 3F).

**Figure 3.**
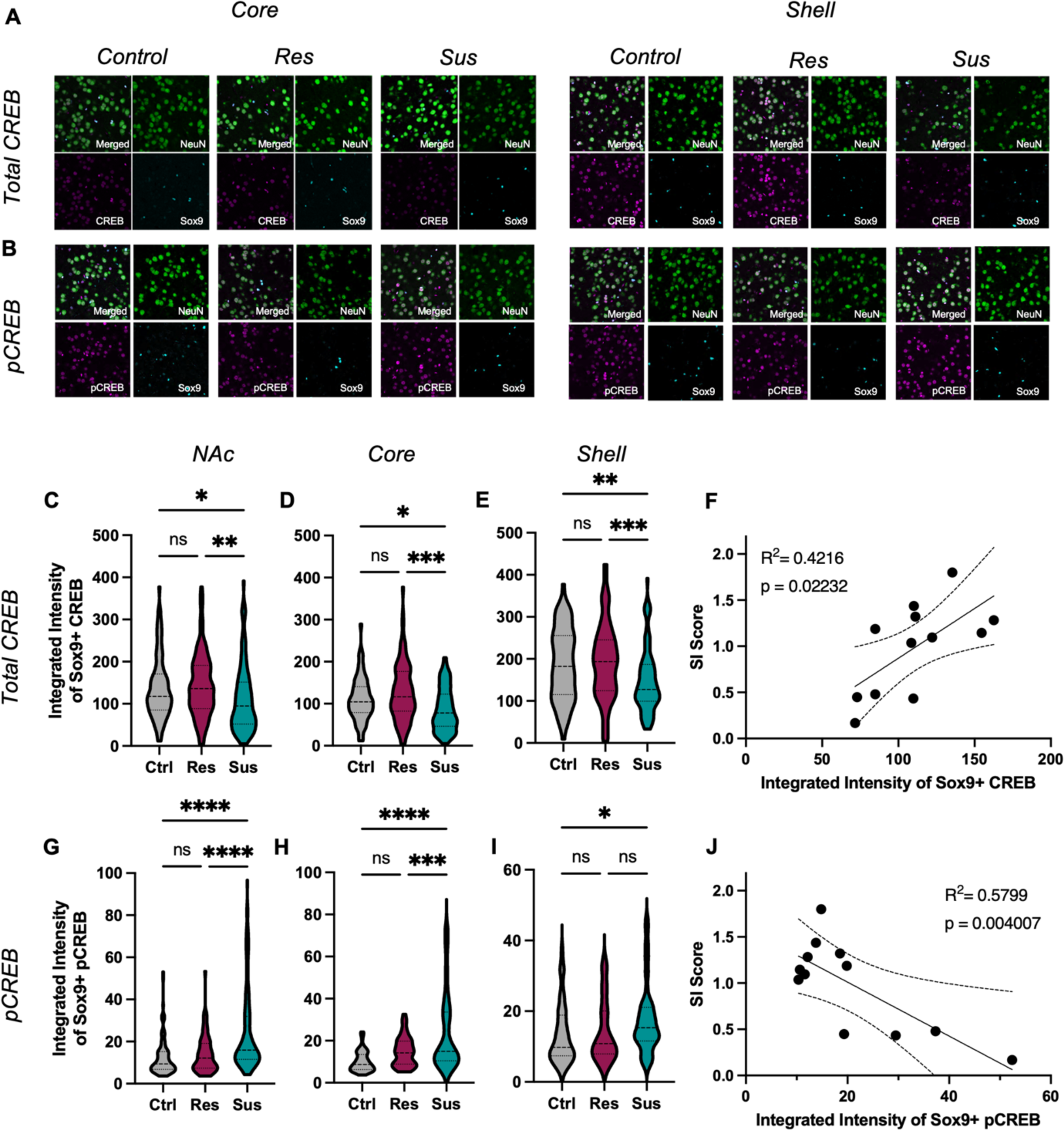
Astrocytic CREB regulation in NAc of susceptible mice. A–B) Representative IHC images for total (A) and pCREB (B) in astrocytes (SOX9, blue) and neurons (NEUN, green) from the core (left) and shell (right) of NAc after CSDS. Across the C) entire NAc, total CREB expression is decreased in Sus astrocytes compared to Res and Ctrl astrocytes (F_(2,337)_ = 6.538, *P* = 0.0016; Tukey post-hoc \**p* = 0.0227; \*\**p* = 0.0019), with no difference between Ctrl and Res astrocytes (Tukey post-hoc *p* = 0.8360). Similar effects were observed in the D) core (F_(2,235)_ = 9.976, *P* = 0.0002; Tukey post-hoc \**p* = 0.027; \*\*\**p* < 0.001) and E) shell F_(2,248)_ = 6.659, *P* = 0.0015; Tukey post-hoc \*\**p* = 0.0157; \*\*\**p* = 0.002). F) Pearson r correlation of individual animal’s mean total astrocytic CREB expression and Interaction Ratio (SI score) reveals a strong positive correlation (r(12) = 0.649, *p* = 0.022, R^2^ = 0.422). G) In contrast, increased expression of the canonical activated form of CREB (pCREB) was observed in Sus compared to Ctrl and Res astrocytes (F_(2,285)_ = 26.72, *P* < 0.0001; Tukey post-hoc \*\*\*\**p* < 0.0001), with no difference between Res and Ctrl astrocytes (Tukey post-hoc *p* = 0.109). This increase in pCREB in Sus astrocytes was observed in H) NAc core (F_(2,178)_ = 23.49, *P* < 0.0011; Tukey post-hoc \*\*\**p* = 0.027; \*\*\*\**p* < 0.001). However, within the shell, increased pCREB in Sus was only observed compared to Ctrl and not Res astrocytes (F_(2,150)_ = 3.760, *P* = 0.0255; Tukey post-hoc \**p* = 0.034). J) Pearson r correlation of individual animal’s mean astrocytic pCREB expression and Interaction Ratio (SI score) reveals a negative correlation (r(12) = -0.762, *p* = 0.004, R^2^ = 0.579). Data represented as mean +/- SEM; n = 5 mice per condition. pCREB: phosphorylated CREB; NAc: nucleus accumbens; CSDS: chronic social defeat stress; Res: resilient; Sus: susceptible; Ctrl: control.

To determine the level of activation of CREB in astrocytes following CSDS we utilized an antibody targeted against phosphorylation of CREB at Ser133, a site canonically associated with increased CREB-mediated transcription [56, 57]. In contrast to total CREB, we found an increase in the integrated intensity of pCREB in Sus astrocytes compared to Ctrl and Res, and no significant effect between Res and Ctrl astrocytes (Fig 3G). Within the NAc core, we again observed increased pCREB in Sus compared to Res and Ctrl, and no significant change in Res compared to Ctrl astrocytes (Fig 3H). A significant increase in pCREB in Sus astrocytes compared to Ctrl was found in the NAc shell, but no difference between Res and Sus or Ctrl astrocytes (Fig 3I). Correlation of the integrated intensity of astrocytic pCREB and SI Ratio revealed a significant negative effect, again suggesting that the change in astrocytic CREB activation is not limited to our categorical stress assignment (Fig 3J). Importantly, and as validation to our above astrocyte findings, similar effects of CSDS on total CREB and pCREB were determined for neurons (NeuN+ cells; (Fig S1A-H) of the same animals, in line with previous reports [50, 58].

### Astrocytic CREB regulates behavioral responses to chronic stress

To study the impact of stress regulation of astrocytic CREB in NAc, we developed AAV vectors to manipulate CREB specifically in astrocytes. We and others have previously utilized viral expression of either a wildtype CREB or a dominant negative CREB mutant termed mCREB (serine to alanine mutation at Ser133) to establish neuronal CREB’s influence on a variety of behaviors [48, 50, 59, 60]. We therefore used these same designs, but with a *GfaABC1D* promotor to target viral-mediated transgene expression selectively to astrocytes in NAc. Immunohistochemistry co-staining with SOX9 and CREB was used to validate astrocyte specificity and revealed that both CREB-GFP+ and mCREB-GFP+ cells only colocalized with SOX9+ cells (Fig 4C), with roughly 70% of NAc SOX9+ astrocytes being GFP+ positive (Fig 4D). To validate that our viruses manipulate CREB expression, a second cohort of mice were injected with unilateral NAc CREB or mCREB AAVs and GfaABC1D-tdTomato AAV (Ctrl-AAV). This injection paradigm allows for the examination of CREB and pCREB expression using a within-subject design to account for individual variability in CREB expression levels. As expected, both CREB and mCREB AAVs resulted in increased expression of total CREB compared to Ctrl-AAV astrocytes (Fig 4E) [61]. Importantly, we only observed increased pCREB integrated intensity in astrocytes expressing CREB-AAV, but not mCREB-AAV (Fig 4F). These data confirm that our viruses selectively manipulate CREB in astrocytes within NAc.

**Figure 4.**
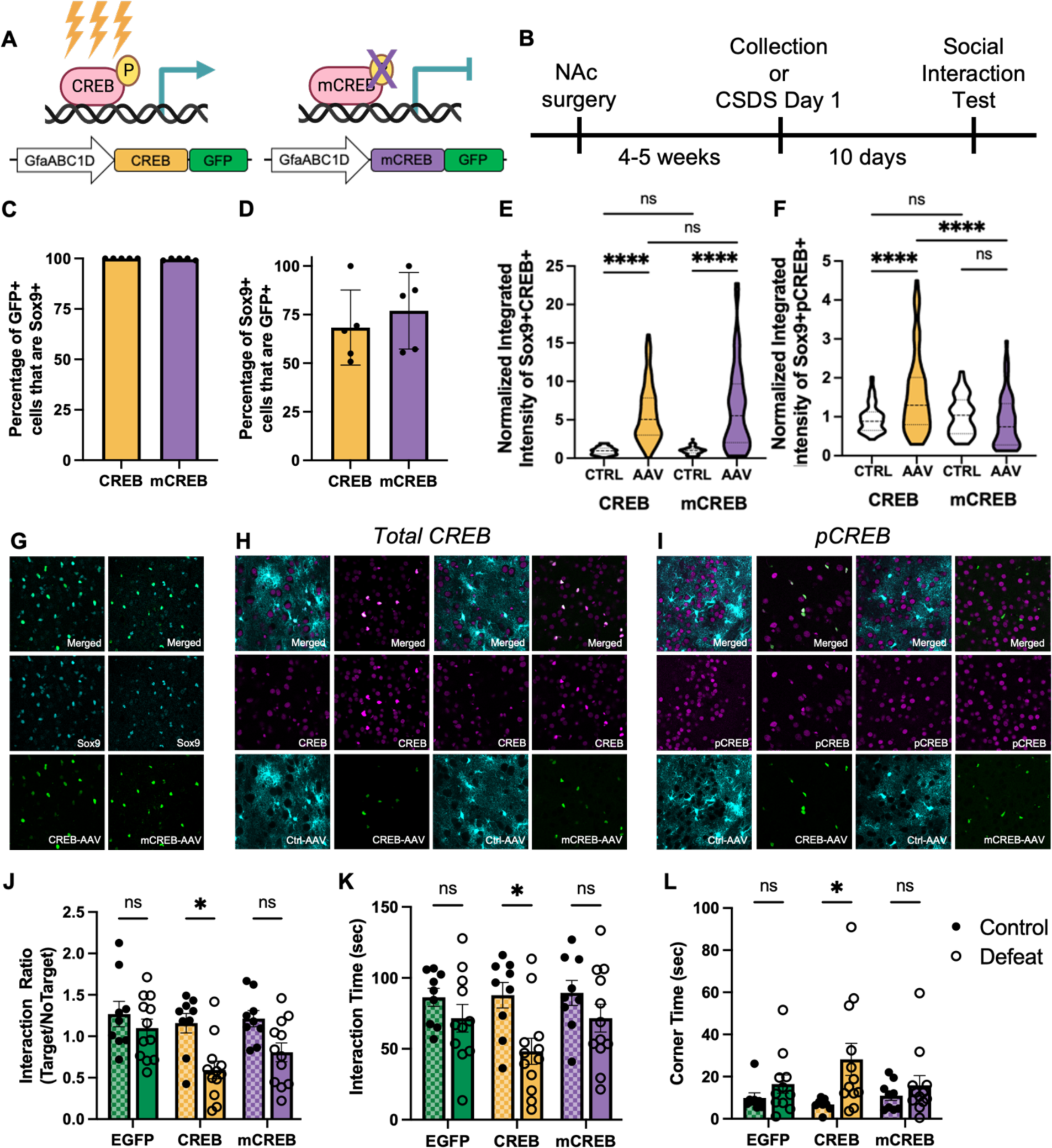
Astrocytic CREB in NAc promotes susceptibility to chronic stress. Cartoon illustrations of A) AAVs to increase CREB activity (left, CREB, yellow) or downregulate CREB activity via expression of a dominant negative mutant (right, mCREB, purple) selectively in NAc astrocytes and B) experimental timeline. C) The percentage of AAV-targeted cells (GFP+) that are astrocytes (SOX9+) demonstrates astrocyte-specific AAV expression. D) Both CREB and mCREB AAVs target roughly 70% of astrocytes within the NAc. E) Increased expression of total CREB was observed with both AAVs, as expected, compared to contralateral Ctrl-AAV expressing astrocytes (F_(3,292)_ = 60.61, *P* < 0.001; Tukey post-hoc \*\*\*\**p* < 0.0001). F) Importantly, only the CREB-AAV increased levels of pCREB (F_(3,246)_, = 14.23 *P* < 0.0001; Tukey post-hoc \*\*\*\**p* < 0.0001), with no difference found in mCREB expressing astrocytes (Tukey post-hoc \*\*\*\**p* < 0.0001). H) Overexpression of CREB selectively in NAc astrocytes decreased the Interaction Ratio following CSDS (two-way ANOVA; main effect of AAV (F_(2,56)_ = 3.46, *P* = 0.0381; main effect of Defeat (F_(1,56)_ = 16.04, *P* = 0.0002) in comparison to both control CREB animals (Tukey post-hoc \**p* = 0.0152) and defeated EGFP animals (Tukey post-hoc *p = 0.0208). I) Decreased Interaction Time (two-way ANOVA; main effect of Defeat (F(1,56) = 9.819, *P* = 0.0027; Tukey post-hoc \**p* = 0.0415) and J) increased time in corners (two-way ANOVA; main effect of Defeat (F(1,56) = 7.431, *P* = 0.0087; Tukey post-hoc \**p* = 0.0450) was also observed in defeated CREB animals compared to control CREB animals. Data represented as mean +/- SEM; for C-F) n = 5 animals per condition; for H-J) n = 9 control and 12 defeat animals per condition).

To determine if astrocytic CREB regulates behavioral responses to CSDS, we used the above AAVs to virally manipulate CREB in NAc astrocytes followed by our 10-day CSDS paradigm. Herein, we included both defeat and viral controls to compensate for any baseline behavioral effects of virally manipulating astrocytic CREB. SI testing was performed 24 hr after the last defeat session. Overexpression of astrocytic CREB was associated with an increased susceptibility phenotype after chronic stress. This effect is particularly striking after categorical assignment ((Fig S1I), with nearly 82% of defeated CREB animals assigned to Sus (9 Sus, 2 Res) compared to 40% in the EGFP control group (4 Sus, 6 Res) or 58% in the mCREB group (7 Sus, 5 Res). Independent of our categorical assignment, overexpression of astrocytic CREB resulted in significantly decreased interaction ratio compared to both control CREB and defeated EGFP animals (Fig 4H). In contrast, no significant difference was found following mCREB astrocytic viral manipulation in comparison to mCREB controls or EGFP defeated animals. Time spent exploring the SI aggressor was also decreased in CREB animals compared to CREB controls, and time spent in the corners was increased (Fig 4I–J). No significant effect was observed for EGFP or mCREB defeated animals. The above data demonstrate that astrocytic CREB biases animals to a susceptible phenotype following CSDS.

## Discussion

The data presented herein are the first to investigate the transcriptional response of astrocytes in the NAc to stress, and one of very few to demonstrate that a transcription factor in adult astrocytes regulates complex behavior. We demonstrate that the NAc astrocytic transcriptome robustly responds to chronic stress, with both a general transcriptional response to chronic stress regardless of phenotype as well as a specific response in resilient versus susceptible astrocytes. Subsequent bioinformatic analysis revealed potential molecular consequences of chronic stress on astrocyte function, including protein stability, cytoskeletal dynamics, and neuronal dysfunction. Given the limited knowledge of astrocytic transcriptional regulators, we additionally performed an Upstream Regulator analysis which deduced CREB as a strong predicted upstream driver, with opposite transcriptional control predicted in Res compared to Sus astrocytes. Thus, the stress-specific response may be mediated by diverging transcriptional regulation in astrocytes, and we indeed found that viral-mediated overexpression of CREB selectively in NAc astrocytes promotes a pro-susceptible phenotypic in response to chronic stress.

Previous work has heavily implicated astrocytes in stress and depression in both rodent models and human MDD subjects [1, 62]. Targeted molecular approaches revealed dysregulated astrocyte gene and protein expression across a variety of brain regions [1]. Within MDD subjects, cell-type deconvolution of bulk tissue RNA-seq data, and single cell RNA-seq approaches, have implicated dysregulation of astrocyte gene expression [4–12, 22]. Investigations of the astrocytic transcriptional and behavioral component have largely been focused on the PFC, hippocampus, or amygdala [6, 8, 11, 17–21]. Here, we performed RNA-seq on whole cell sorted astrocytes after chronic stress. We chose to focus on the NAc given its important role in motivation, reward, and emotion-related behaviors and evidence for astrocytic regulation by stress or MDD from bulk RNA-seq studies in mice and humans, respectively [22, 26]. We found a robust transcriptional response to stress in both Res and Sus astrocytes, including a set of gene changes unique to Res, which highlights that resilience to stress is an active biological process as seen previously for neurons [9, 26, 63]. Examination of statistically significant DEGs revealed little overlap between Res and Sus astrocytes; however, threshold-free RRHO2 analysis revealed concordant patterns of gene expression. We observed that Res astrocytes generally demonstrated a larger Log_2_FC compared to Sus astrocytes. Therefore, the lack of overlap may simply be due to a Sus transcriptional response that does not reach statistical significance. On the other hand, our results may reflect both a general astrocytic transcriptional response to stress, as well as phenotype-specific transcriptomes. Importantly, comparison to bulk RNA-seq from human MDD patients revealed concordant gene expression in both Sus and Res astrocytes, although with a stronger overlap between Sus and MDD. This finding additionally suggests both a general and specific astrocyte transcriptional response to stress and depression.

Despite an increase in sequencing studies directed at astrocyte populations, we know surprisingly little about astrocytic transcriptional regulators. Bioinformatic analysis of our data highlighted several predicted upstream regulators of the astrocyte transcriptional response to CSDS, including some previously implicated in social behavior, stress, or depression [37, 52–55]. Noteworthy, both CD38 and TCF7L2 were found to be important for proper neurodevelopment associated with social behavior, but within the PFC [52, 53]. While these regulators were associated with younger postnatal ages, a recent study by Huang et *al*. demonstrated that global loss of astrocytic Nuclear Factor-IA (NFIA), a well-known transcription factor associated with astrogliogenesis during neurodevelopment, in adult animals resulted in hippocampus-specific effects on transcription and behavior [64]. In the evolving field of astrocyte biology, there is a growing need to investigate the regional, temporal, and contextual role of astrocyte transcriptional regulation.

Our analysis additionally revealed CREB as a strongly predicted upstream regulator, but with opposing activation states in Res compared to Sus astrocytes. Previous work has also implicated CREB as a transcriptional regulator in cultured astrocytes [33–36]. We confirmed this bioinformatic result with examination of the levels of total and activated CREB (pCREB) in astrocytes at the protein level. We found increased expression of activated CREB in Sus NAc astrocytes following CSDS, consistent with the Upstream Regulator analysis. Furthermore, the levels of total and pCREB correlated with an individual animal’s SI score, albeit in opposite directions, indicating that our results are not merely due to nominal discrete groupings. Instead, this indicates that CREB expression influences responses to stress on a continuous scale. Concomitantly, we found the same results in neuronal populations from the same animals, in line with previous work and in support of our astrocytic results [48–50]. Neuronal CREB displays a region-specific influence on stress and depression, wherein a pro-resilient effect is associated with neuronal CREB expression in the PFC and hippocampus, but a pro-susceptible effect in NAc [65]. Astrocyte regional heterogeneity is well documented; thus, it is reasonable to assume a region-specific influence of astrocytic CREB [64, 66–69]. Nevertheless, future work is needed to determine if astrocytic CREB also displays region-specific influence on behavioral responses to stress.

How transcriptional regulation in astrocytes contributes to complex behavior is an exciting and emerging field. We examined, for the first time, the role of astrocytic CREB in regulating stress responses and demonstrated that astrocytic CREB induces a pro-susceptibility phenotype. This effect was particularly prominent following categorical assignment to Res or Sus categories, with over 80% of animals assigned to Sus. Outside of discrete grouping, this effect was still observed, with a significant reduction in SI Ratio. An important limitation to our study is the inclusion of only males. While sex differences in CREB’s association with MDD are inconclusive, there are known sex differences in both human MDD and rodent models [22, 42, 70–72]. Future studies are needed to determine the sex-specific astrocytic transcriptional response to chronic stress and how astrocytic CREB may regulate depressive-like behaviors in a sex-specific manner. Interestingly, we did not observe a change in SI ratio after viral expression of mCREB, a dominant negative mutant, in NAc astrocytes. This was surprising, as previous work in neurons demonstrates that this construct induces opposite molecular and behavioral changes compared to wildtype CREB [58, 59, 61]. However, the exact molecular mechanisms by which astrocytic CREB induces transcription are not fully understood, and cell culture studies report contradictory results [34–36]. It is possible that compensatory molecular mechanisms may be sufficient to overcome the effects of virally-expressed mCREB.

To conclude, we demonstrate that the astrocyte transcriptome within the NAc robustly responds to CSDS in both resilient and susceptible astrocytes. We furthermore demonstrate that transcriptional regulation in astrocytes mediates depressive-like behaviors, as viral overexpression of CREB selectively in NAc astrocytes biased animals towards a susceptible phenotype. Our data strongly support the increased attention on astrocytic responses in stress and depression research and highlight the importance of better understanding transcriptional regulation in astrocytes which may reveal yet unknown molecular mechanisms underlying neuropsychiatric disorders which can be targeted with novel therapeutics.

## Acknowledgements

This work was supported by funding from the NIH (R01MH051399 and R01MH129306 to E.J.N.). The authors would like to thank Katherine Beach, Catherine McManus, Kyra Schmidt, Nathalia Pulido, and Ezekiell Mouson for animal husbandry.

## Disclosures

The authors declare no competing financial interests.

## Data Availability

All RNA-seq data reported in this study will be deposited in the Gene Expression Omnibus.

---

**Supplemental Figure 1.**
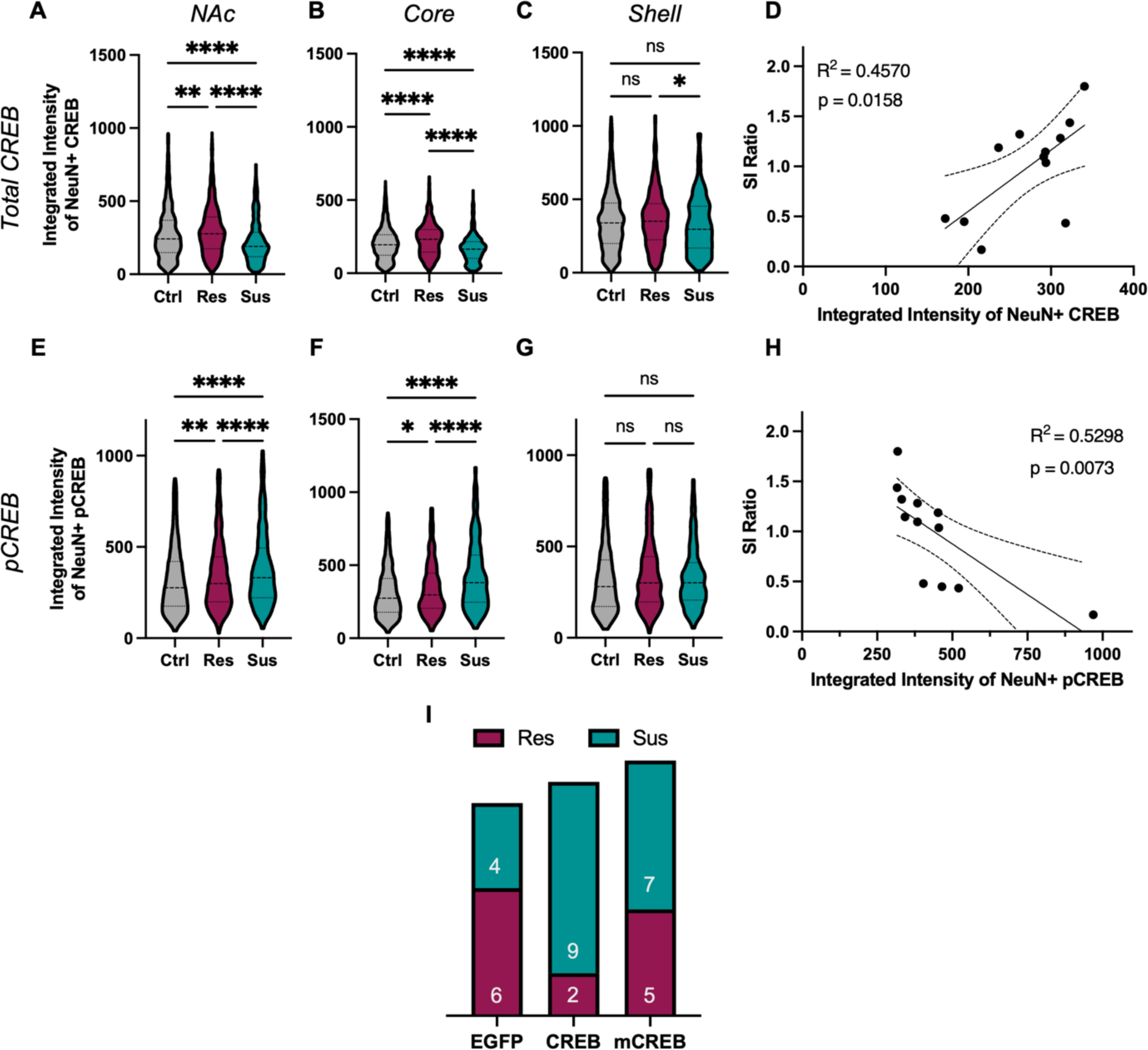
Across the A) entire NAc, total CREB expression was decreased in Sus neurons compared to Res and Ctrl neurons, and increased in Res neurons compared to between Ctrl (F_(2,3275)_ = 50.60, *P* < 0.0001; Tukey post-hoc \*\**p* = 0.0016; \*\*\*\**p* < 0.0001). Similar effects were observed in the B) core (F_(2,1717)_ = 51.68, *P* < 0.0001; Tukey post-hoc \*\*\*\**p* < 0.001). C) In the shell, total CREB expression was only decreased in Sus neurons compared to Res, with no significant different compared to Ctrl (F_(2,1568)_ = 4.271, *P* = 0.0141; Tukey post-hoc \**p* = 0.012). D) Pearson r correlation of individual animal’s mean total neuronal CREB expression and Interaction Ratio (SI score) revealed a strong positive correlation (r(12) = .676, *p* = 0.0158, R^2^ = 0.4570). E) In contrast, increased expression of pCREB was observed in Sus and Res compared to Ctrl neurons (F_(2,3762)_ = 36.28, *P* < 0.0001; Tukey post-hoc ** *p* = 0.0013, \*\*\*\**p* < 0.0001) in total NAc. This increase in pCREB in Sus neurons was observed in F) NAc core (F_(2,1951)_ = 56.72, *P* < 0.0001; Tukey post-hoc \*\*\*\**p* < 0.001), but not in the G) shell (F_(2,1833)_ = 2.782, *P* = 0.0622). H) Pearson r correlation of individual animal’s mean neuronal pCREB expression and Interaction Ratio (SI score) revealed a negative correlation (r(12) = -0.728, *p* = 0.0073, R^2^ = 0.5298). I) Phenotype assignment for defeated animals following viral-mediated astrocytic CREB manipulation and CSDS. Data represented as mean +/- SEM; n = 5 mice per condition. pCREB: phosphorylated CREB; NAc: nucleus accumbens; CSDS: chronic social defeat stress; Res: resilient; Sus: susceptible; Ctrl: control.

## Notes

### Competing Interest Statement

The authors have declared no competing interest.

